# A synthesis of senescence predictions for indeterminate growth, and support from multiple tests in wild lake trout

**DOI:** 10.1101/2021.10.05.463025

**Authors:** Craig F. Purchase, Anna C. Rooke, Michael J. Gaudry, Jason R. Treberg, Elizabeth A. Mittell, Michael B. Morrissey, Michael D. Rennie

## Abstract

Senescence, or the deterioration of functionality with age, varies widely across taxa in pattern and rate. Insights into why and how this variation occurs are hindered by the predominance of lab-focused research on short-lived model species with determinate growth. We synthesize evolutionary theories of senescence, highlight key information gaps, and clarify predictions for species with low mortality and variable degrees of indeterminate growth. Lake trout are an ideal species to evaluate predictions in the wild. We monitored individual males from two populations (1976-2017) longitudinally for changes in adult mortality (actuarial senescence) and body condition (proxy for energy balance). A cross-sectional approach (2017) compared young (ages 4-10 years) and old (18-37 years) adults for (1) phenotypic performance in body condition, and semen quality - which is related to fertility under sperm competition (reproductive senescence), and (2) relative telomere length (potential proxy for cellular senescence). Adult growth in these particular populations is constrained by a simplified food web, and our data support predictions of negligible senescence when maximum size is only slightly larger than maturation size. Negative senescence (aka reverse senescence) may occur in other lake trout populations where diet shifts allow maximum sizes to be much larger than maturation size.

## 1. INTRODUCTION

Senescence is a decline in individual biological function with age, and is typically quantified as an increase in adult mortality rate or reduced ‘fertility’ [1], but can be applied to any decline in phenotypic performance. Tremendous variability exists among species in the shape (direction) and speed (rate) of senescence [2–5], and many authors seek to explain such patterns [e.g., 3, 6, 7]. The contention that the strength of selection declines with age is a common explanation of senescence [8]. The premise being that few individuals reach old age, and many have already reproduced at younger ages, therefore, selection cannot remove problematic traits that arise only at old age. An hypothesis that “low adult death rates should be associated with low rates of senescence, and high adult death rates with high rates of senescence” [9], has empirical support. However, the nuances of the hypothesis and its predictions are debated [6, 10, 11]. Relative adult to juvenile mortality appears critical [6], but asymmetry between parent and offspring [7] can differ widely between determinate and indeterminate growers and generalizations can be problematic. An example with bivalves provides a useful illustration [see 6, page 527], which would also apply to most fishes.

Our manuscript has three primary goals: 1) synthesize existing senescence theories, showing the importance of growth pattern, and highlight types of data needed to fill key voids, 2) introduce lake trout (*Salvelinus namaycush*) as an ideal species to address senescence in the wild, 3) present a case study of two lake trout populations with exceptional monitoring.

### 1.1 EVOLUTIONARY THEORIES OF SENESCENCE

Attempts to explain senescence are challenged by inconsistencies in terminology and in the hierarchy of how theories are grouped. Complicating things further, the major theories of senescence [7] are not mutually exclusive, and create similar predictions but for different reasons. Our interpretation (Figure 1) represents a modification from Maklakov and Chapman [8; their Figure 2]. The mutational accumulation theory (MAT, Figure 1), posits [12] that individuals senesce due to the accumulation of deleterious mutations through their lifetime, such that senescence is strictly maladaptive. Other theories (Figure 1) consider the notion of fitness optimization or life history tradeoffs, whereby declining performance with age may result from increased performance while young. The antagonistic pleiotropy hypothesis (APH, [9]) suggests senescence occurs when certain genes have positive effects in early life but negative effects later. The disposable soma hypothesis (DSH) proposes [13] energy allocated in reproduction is unavailable to maintain soma, resulting in deterioration. Many present APH and DSH as distinct, but we consider DSH to be a version of APH (Figure 1). More recently, optimization of function has been proposed; appearing as developmental function theory (DFT, [8]) and hyperfunction [14]. Conceptually this is similar to DSH but the proposed mechanism varies, being energy allocation for DSH (a tradeoff) and hyperfunctioning genes that lead to excessive biosynthesis and molecular turnover in mature individuals for DFT (which unlike [8] we consider as a putative constraint [sensu 15] – as opposed to a plasticity enabled tradeoff; Figure 1). How DFT might apply to indeterminate growers is unclear, as development never stops.

**Figure 1:**
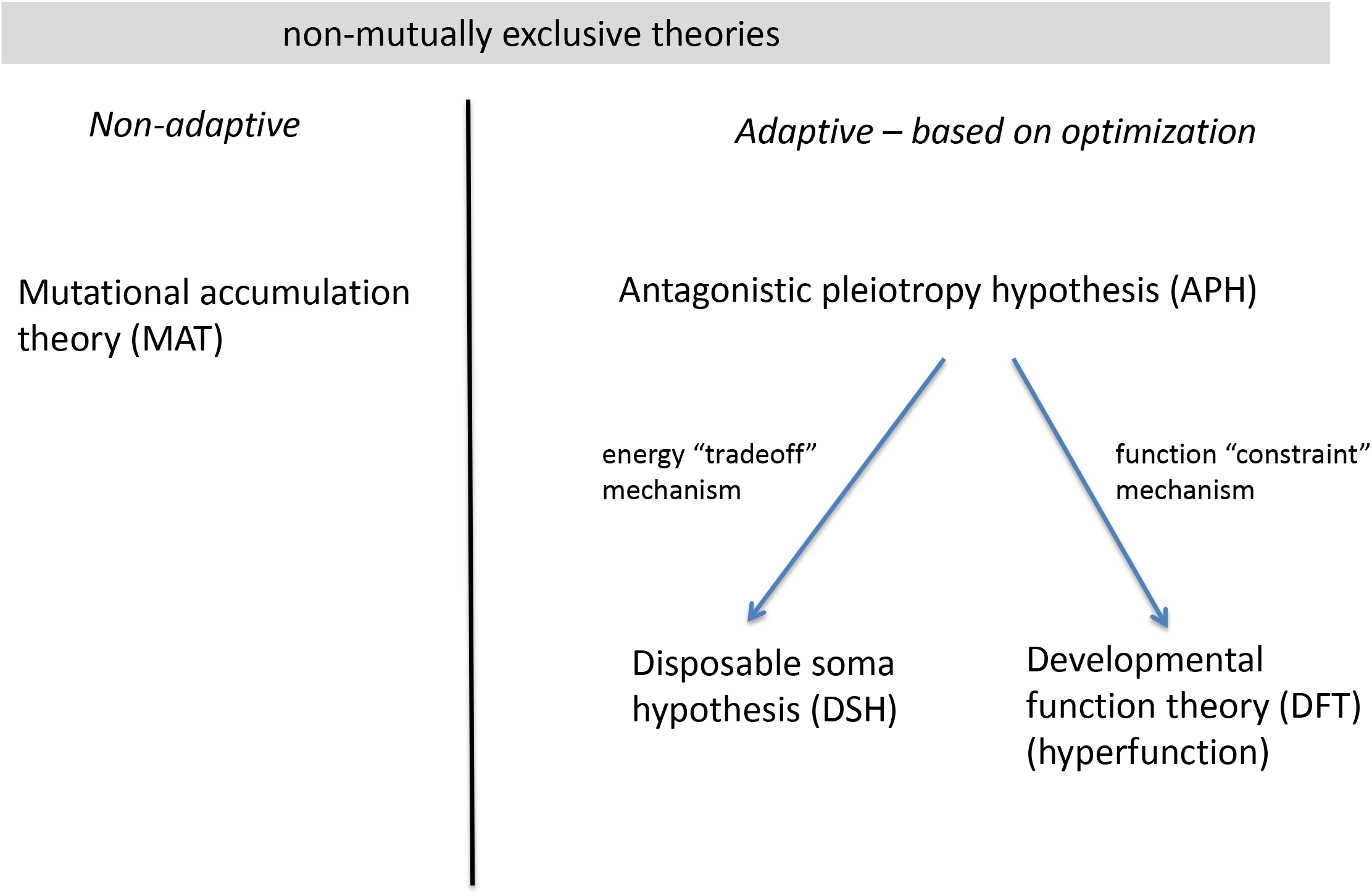
Our hierarchical conceptualization of the main evolutionary theories of senescence. Modified and expanded form Maklakov and Chapman [9].

### 1.2 ATYPICAL PATTERNS OF SENESCENCE

Most empirical work on senescence has been framed in support of DSH [e.g, 8, 15, 16]. However, there has been recent questioning of this [8, 15, 17], and new research that addresses some key gaps may be revealing. Examining unusual patterns of senescence [3, 7] may help illuminate why and how it occurs ([5]; Figure 1). Negligible senescence describes species with little or no deterioration with age [2, 18, 19], while negative (reverse) senescence [20] may occur when biological function increases with age. The tenet of this argument is that in all species, mature individuals have offspring that are smaller than themselves. As offspring grow, their ability to reproduce increases and their probability of mortality can decline. In species with determinate growth, this pattern stops at maturity. Indeterminate growers however continue to increase in size after maturity. If mortality declines and fertility increases with size (age), then there is increased selection against senescence in indeterminate versus determinate growers.

Across different conditions, an optimization model [20] concludes that the intrinsic growth pattern (determinate vs indeterminate) influences the shape (direction) of senescence, while mortality determines its rate. Predictions can be summarized as: (1) senescent conditions (classical ageing) occur when the size at maturity is close to the maximum size (determinate growth) with little scope for increasing fertility with age (e.g., mammals, birds, insects); (2) negative (reverse) senescence should occur when size at maturity is much less than maximum size (some indeterminate growers), and reproductive capacity increases with size; (3) negligible senescence (little ageing) is an arbitrary middle ground along this continuum and should occur when size at maturity is somewhat less than maximum size, but reproductive capacity increases with size (age). Support for this framework appears in a recent review [7].

### 1.3 DESIRABLE STUDY SYSTEMS TO FILL KEY VOIDS

Studies of senescence are heavily skewed towards a narrow range of conditions. A synthesis of the repeated calls [e.g., 16] to address knowledge gaps includes:

(1) A critical need for research focusing on species with indeterminate growth [1, 20–22], for example in certain plants [23], reptiles [24] and fishes [18, 19]. Most work on senescence has considered determinate growers (mammals, birds, insects), which may bias our view of ageing.

(2) A requirement to examine senescence in wild populations [1, 25, 26], which better encapsulate natural processes and influences of potential environmental covariates on senescence. Laboratory studies of model organisms lack this relevance.

(3) Research that combines both longitudinal and cross-sectional comparisons of age is valuable [e.g., 27]. Comparisons in fitness-related traits can be made among age classes (cross-sectional) or by following individuals through time (longitudinal; [23]). Because long-lived individuals may have inherent higher quality, their presence may bias cross-sectional comparisons, making longitudinal studies a desired approach [26, 28]. However, longitudinal studies are subject to other confounding variables (e.g., directional environmental change), and it can take decades to track new metrics if following future cohorts. Thus, studies reporting consistent conclusions across combined approaches may provide more robust tests of hypotheses.

(4) Examinations of wild populations not subject to confounding variables [29], such as immigration/emigration (which may influence estimates of adult mortality), anthropogenic effects (e.g., recent changes in mortality adding novel selective pressure), and adult diet shifts with increasing body size, which can have dramatic influence on reproduction (e.g., gape limited carnivorous reptiles shift diet and are a problem, filter feeding bivalves are not).

(5) Research using recognized cellular indices associated with senescence, like relative telomere length [30] and the influence of reactive oxidative species and their potential for oxidative stress or cellular damage [29] are needed [26, 31], particularly in wild ectotherms. Evolutionary literature on senescence ponders what happens (patterns), why it happens (or does not), but rarely addresses how it happens [8, 28, 31, 32]. Laboratory and model organism-based studies on the biochemical mechanisms, or at least correlates associated with aging and senescence, provide a framework that can be applied to study senescence in the wild.

(6) Research focusing on reproductive senescence [24, 26, 33]. Most studies [26] of senescence quantify it as change in adult mortality rates (actuarial senescence), yet invoking mortality as an explanation is circular [26, 34] being both a cause and consequence of senescence. Measures of reproductive senescence are free of this problem, as are other phenotypic traits.

(7) Senescence research that considers male individuals. Females have been the historical focus for senescence research [32, 33], but in most cases, males should senesce faster [8, 32, 35–39] thus offering larger effect sizes and greater power to answer key questions. This is most pronounced in species with intense sexual selection [36, 40] as increased reproductive effort may come at a cost to tissue maintenance, and mortality can be consistently higher on males due to conspicuous displays.

#### • The special problem of sperm senescence

Reproductive senescence includes senescence on the adult individual (such as ability to attract a mate), but additionally on gametes [31, 41–43]. Gamete senescence affects the fitness of the individual, but also its mate and offspring [43, 44]. However, separating effects of the parent, gamete, and offspring is difficult, especially in internal fertilizers. Egg senescence is rarely measured [33], but sperm senescence is gaining interest [43, 45]. Sperm senescence can be considered in two phases [42, 45]: pre-meiotic (how the age of the male influences sperm) and post-meiotic (both before and after ejaculation). Sperm are particularly vulnerable to oxidative damage [31], and the male mutational bias [42, 46], has led to interest in human fertility and paternal effects. Male fitness is a function of mating opportunities, sperm performance and offspring viability [33, 44], which can be separated under experimental conditions [e.g., 47, 48]. Older males generally produce sperm with reduced fertilization ability [27, 29, 33] and lead higher rates of developmental abnormalities among offspring [29].

## 2. LAKE TROUT

### Desirable attributes

Lake trout present an ideal indeterminate growth model for studies of senescence in nature, with low adult mortality being a key attribute. They inhabit the hypolimnion of lakes [49], where there are functionally no predators on adults (contrasts greatly to marine predation on anadromous salmonids) and spawn on lake shoals at night [49, 50], where they are not exposed to terrestrial predators (unlike stream spawning salmonids).

Reproductive quality and investment can be accurately estimated from gametes. Lake trout do not typically migrate to spawn, show few secondary sexual characteristics, no sexual dimorphism, have no energetically costly courtship, and provide no parental care [49, 50]. Fertility increases with size (age), as larger females produce more eggs. Males do not compete for territories [49, 50], but post-ejaculatory sexual selection [44] occurs due to sperm competition [50]. Larger (older) fish generally produce more sperm, and thus would gain paternity advantages (fertility) under a fair raffle system [51].

Variation in maximum body size across populations (variable realization of indeterminate growth) may be useful for testing predictions of negligible and negative senescence [20] within the same species. Lake trout are amongst the largest members of the Salmonidae family, but maximum body size varies greatly as a function of prey availability [52, 53]. Thousands of populations vary in life history traits that influence their fitness [54]. Inter-population comparisons could exploit environmental variation (something senescence literature has been asking for [e.g., 15, 16, 26, 28, 32]) in variables such as growing season, prey resources, and juvenile predators.

### Support for theories of ageing

If senescence is optimized (Figure 1) between fitness benefits early in life at a cost to either hyperfunctioning genes (DFT) or somatic maintenance (DSH), then selection against a decline in performance with age is predicted to be relatively high in lake trout, as fitness potential increases dramatically with size (age), given adult mortality rates decline while fertility increases. We are unaware of any published data that can shed specific light on DFT in lake trout. However, low allocation in reproduction is predicted to plastically tradeoff with high investment in somatic maintenance under DSH [55]. Possibly supporting this, lake trout have relatively low secondary sexual characteristics/migration/courtship/fecundity (resulting in low annual reproductive effort) and a predictably high incidence of iteroparity [49]. Perhaps consequently, they can live to ages of >60 years [56], making them among the longest lived fishes, vertebrates, and animals on the planet. Using a variety of approaches, we sought to directly test the hypothesis that wild lake trout show little or no senescence [20].

### Case study of two populations

Our study populations have several additional attributes making them valuable for testing hypotheses of senescence in the wild. Many potentially confounding variables can be ruled out, as the lakes are located at the IISD Experimental Lakes Area (Ontario, Canada), where recreational fishing is prohibited and there is no unquantifiable directed anthropogenic activity. Annual mark-recapture studies have been ongoing since 1976, enabling long-term monitoring of individuals. The lakes are very small (see methods) and all adults of various ages within a population experience similar environmental conditions. There are no piscivorous predators (except lake trout), adult trout are too large to be taken by loons (*Gavia immer*), but might occasionally be prey to otters (*Lontra canadensis*). Adult mortality is thus very low, whereas mortality of small juveniles is likely relatively high [sensu 6]. The lakes are connected in their surrounding watershed by very small streams, effectively eliminating immigration/emigration for this hypolimnetic species. Due to a simplified food-web [52, 57] adult trout in these two lakes do not switch diet as they age, and gain little body size after maturity (Figure 2a, and published growth curves [57]). This is critically important, as diet is known to affect gamete quality in fishes [e.g., 58] and would bias age (size) comparisons in most systems. Sampling over the course of 40+ years has shown that young and old adult male lake trout co-occur on the spawning shoals at the same time (Rennie, unpublished), thus our comparisons of age are not confounded by differential spawn timing.

**Figure 2:**
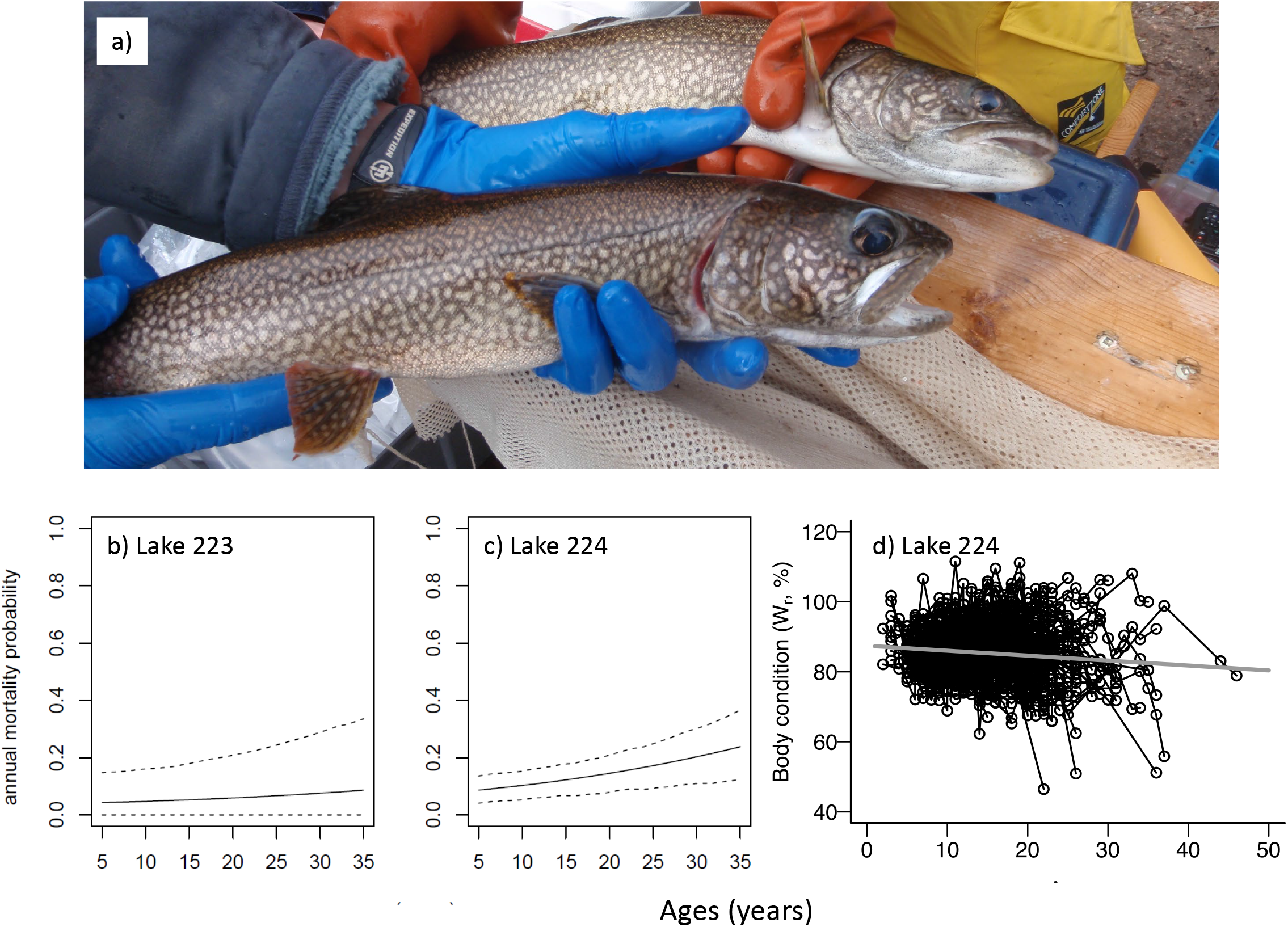
Longitudinal data from individually tagged adult male lake trout. a) adults of age 9-years (back) and 37-years (front); b) and c) annual mortality probability as a function of continuous age in Lake 223 and Lake 224, respectively. The solid line is the mean predicted probability, and the dashed lines are the 95% credible intervals; d) body condition relative to age from Lake 224. Each black line is a resampled fish (minimum six times) during October, 1976-2017.

## 3. METHODS

In polyandrous mating systems like lake trout, male “fertility” is influenced by the ability to achieve fertilizations under sperm competition [33, 44], a key component [45] being sperm swimming performance. We thus quantified male “fertility” by measuring sperm traits that predict paternity. We also measured adult mortality estimates, body condition as a surrogate for general health [59, 60], and relative telomere length as a cellular-level marker of senescence [61–63]. Our study thus combines actuarial senescence, phenotypic measures of bodily function with age (including reproductive senescence), along with a potential biochemical senescence marker, providing a more holistic approach others have highlighted as being needed [e.g., 8].

### 3.1. LONGITIDUTINAL STUDY

At first capture, fish were tagged, measured (total length, mass) and sexed, with the leading fin ray of a pectoral fin removed for ageing [64]. Recaptures in subsequent years used tag identification to assign age. Fish over the entire duration of monitoring in Lake 224 (27.3 ha, 1976–2017) were used, while from Lake 223 (26.4 ha) we restricted data to 1990-2017, to exclude the potential influence of an historical acidification experiment [57] – too few samples remained to track condition in Lake 223.

#### (A) Actuarial Senescence

We estimated annual individual recapture and mortality probabilities using all adult males with known ages (Lake 223 = 385, Lake 224 = 422). To test for changes in adult mortality with age, we fitted a Cormack Jolly Seber model with a Bayesian framework (see Supplemental Methods). Recapture and mortality probabilities were modelled as logistic regression functions of age, which was treated as a continuous variable.

#### (B) Phenotypic performance senescence – body condition

Length-based body condition was estimated as a percentage of standard weight [65]. Fish from Lake 224 that were recaptured at least 6 times during their adult life were used to determine if condition declined with age, and were analyzed with a mixed effects modelling framework (Supplemental Methods). Condition was evaluated as a function of fish age (fixed effect), and repeated measures on the same individuals (random slope), and the year sampled (random intercept).

### 3.2. CROSS-SECTIONAL STUDY

#### (A) Fish collection

We collected fish on spawning shoals at night from 11 to 16 October 2017 and sampled the next morning following previous procedures [66]. Ages of recaptured fish were determined in the field by cross-referencing a database of tag IDs. Younger adult trout were more abundant than older individuals. To avoid potential confounding variables associated with date of sampling (e.g., weather, transport time to laboratory), we grouped fish as either being young (ages 4–10) or old (18–37) and processed them in a ‘group design’ (i.e., the same number of young and old fish were sampled each day). We analyzed 15 groups in each lake (60 total; Supplemental Methods).

#### (B) Sample collection

Eggs were extruded from one female each day and later separated from ovarian fluid through a fine meshed net [67], which was used in sperm swimming performance trials [68], to avoid neutral sperm swimming environments when post-ejaculatory sexual selection occurs [29]. From each male, blood was taken from the caudal peduncle and semen was expressed by gentle abdominal massage. All samples were immediately immersed in ice, and transported to the lab for further processing (completed within 8 hours of collection).

Aliquots of blood and semen were removed from ice and centrifuged (5000 × g at ~15°C for 5 mins). Prior to freezing in liquid nitrogen, plasma was separated from blood cells. A separate semen aliquot was centrifuged in hematocrit tubes, and spermatocrit was computed [69]. This correlates with semen sperm density and often varies within individuals through a spawning season [e.g., 70].

#### (C) Sperm swimming performance

Details (Supplemental Methods) closely followed Purchase & Rooke [67]. Four technical replicates of sperm activation were obtained for each fish. We were able to get useful data within 6 s of sperm/media mixing. Videos of swimming sperm were analyzed in 0.5 s increments using open source software [71]. We used sperm curvilinear swimming velocity (VCL; μm/s) as a metric of male fertility, as it has been repeatedly shown to be correlated to paternity under sperm competition [72].

#### (D) Relative telomere length

We measured relative telomere length from DNA recovered from red blood cells and sperm pellets using a qPCR-based approach that produces a telomere repeat (T) to single gene (S) copy number ratio (T/S). The assay was performed with two single copy genes, *orexin* (*Ox*) and *follicle stimulating hormone beta subunit* (*FSH*), to verify consistency of T/S ratios (Supplemental Methods). Both genes (*Ox* and *FSH*) garnered congruent relative T/S ratios (Pearson’s correlation; blood: *r* = 0.67, *P* < 0.0001, sperm: *r* = 0.72, *P* < 0.0001), thus only the results of *Ox* are presented.

#### (E) 2017 cross-sectional statistical analyses

Body condition, spermatocrit, and relative telomere length were evaluated as a function of fish age (young vs. old) crossed with lake of origin. Sperm swimming declines rapidly after activation, with most successful fertilizations occurring in the few seconds after release. As such, we quantified sperm swimming using two approaches. First, to assess maximum swimming speed, we measured sperm at 6 s post-activation as a function of fish age (continuous variable: 4–37 years) crossed with lake, including tag ID (random intercept) to account for the four technical replicates per male. We also tested for changes in sperm swimming speed over time post-activation (continuous: 6–30 s) crossed with age (young vs. old) and lake. Tag ID (random slope and intercept) and technical replicate (random slope and intercept) were included. In all cross-sectional analyses the interaction between age and lake was not significant (*P* > 0.23), indicating that the effect of age was similar in both populations. We removed these non-significant interactions prior to reporting final model results.

## 4. RESULTS

### ACTUARIAL SENESCENCE

Annual mortality probability estimates of adult male lake trout were low (< 0.20) across all ages in both lakes, and suggest a modest increase with age (Figure 2b, c). This effect of age was clearer in Lake 224 compared to Lake 223 (99.8% and 80.5% of the posterior distributions of the slope parameter were positive, respectively).

### PHENOTYPIC PERFORMANCE SENESCENCE

#### Longitudinal condition

Accounting for random individual (194 fish, 1608 observations) and annual variation, there was a significant change in adult body condition with age in Lake 224 (t_216.2_ = −2.6, *P* = 0.009; Figure 2d). The rate of decline was negligible at 1.4 units per decade, which is well within the variation among fish and years (most observations between 70-105 units).

#### Cross-sectional condition (2017)

Overall mean body condition in October 2017 was 83.4 +/− 0.9%, similar to historical records (Figure 2d). Body condition was similar in both lakes (Lake 224 – Lake 223 means +/− SE: 0.053 +/− 1.79%, t_55_ = 0.03, *P* = 0.976) and there was no difference between young and old trout (old – young means +/− SE: −0.146 +/− 1.79%, t_55_ = −0.082, *P* = 0.935; Figure 3a).

**Figure 3:**
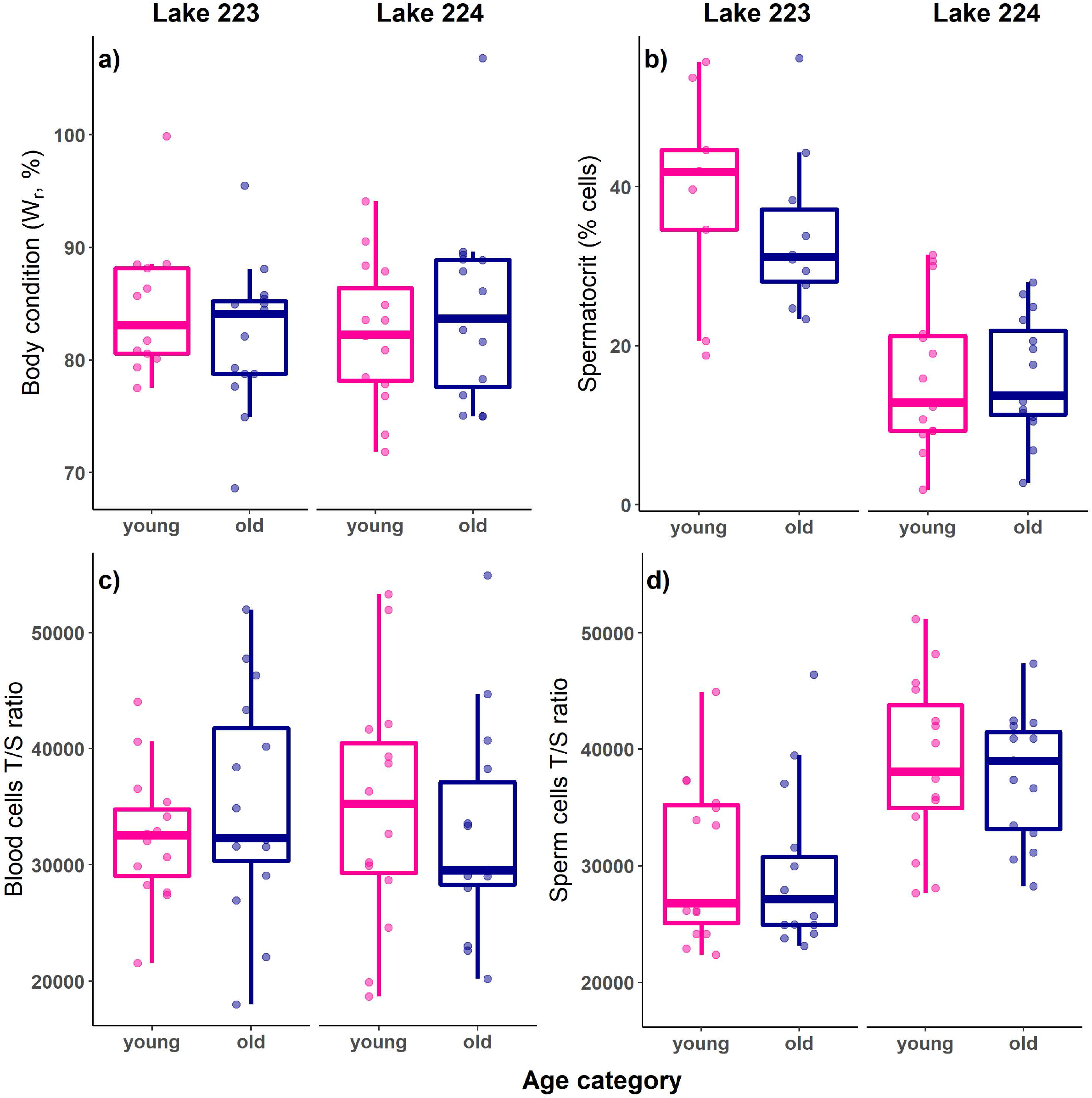
Phenotypic measures of young (pink: 4–10 years, n = 30) and old (blue: 18–37 years, n = 30) adult male lake trout from Lake 223 (n = 30) and Lake 224 (n = 30) in October 2017. Each point represents an individual trout. (a) Body condition, (b) spermatocrit, relative telomere length of (c) red blood cells, and (d) sperm cells. Telomere data presented using Ox reference gene, points represent average of three technical replicates per individual.

#### Cross-sectional semen quality (2017)

Although spermatocrit was significantly higher in Lake 223 in 2017 (Lake 224 – Lake 223: −0.203 +/− 0.028%, *t*_46.0_ = −7.21, *P* <0.0001), there was no difference between young and old fish (old – young: −0.019 +/− 0.028%, *t*_46.0_ = −0.70, *P* = 0.49; Figure 3b). Swimming speed at 6 s post-activation was not significantly different between lakes (Lake 224 – Lake 223: −15.0 +/− 8.1 μm/s, t_56.9_ = −1.84, *P* = 0.071), and was not related to fish age (0.71 +/− 0.45 μm/s per year, t_56.9_ = 1.58, *P* = 0.12; Figure 4a). The rate of decline in swimming speed over time post-activation was faster in Lake 223 (rate difference, Lake 224 – Lake 223: 0.97 +/− 0.37 μm/s per second post activation, t_57.0_ = 2.65, *P* = 0.01); however, there was no difference between young and old trout (rate difference, old – young: −0.43 +/− 0.37 μm/s per second post activation, t_57.0_ = −1.18, *P* = 0.24; Figure 4b).

**Figure 4:**
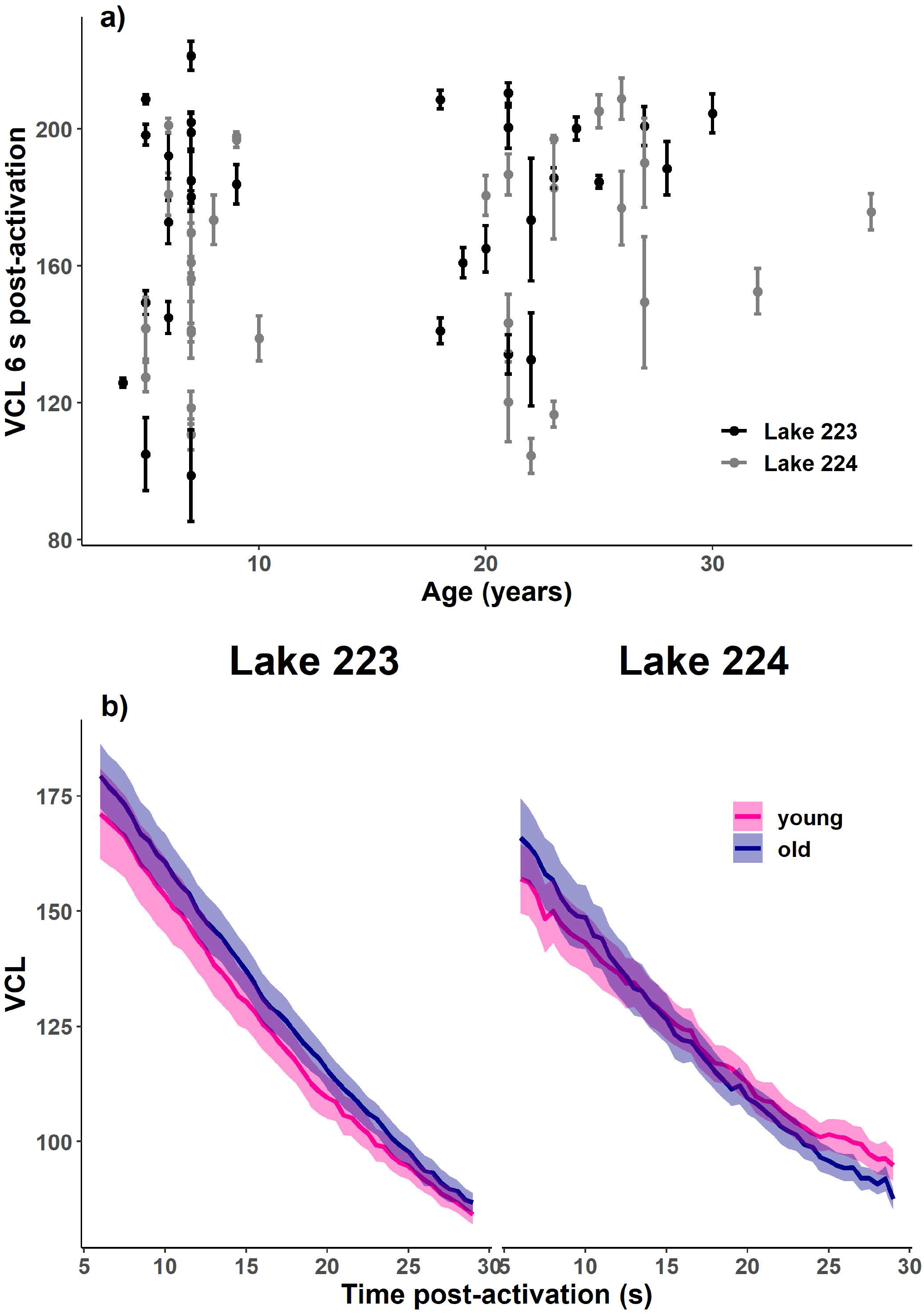
Sperm swimming speed (μm/s) of lake trout in October 2017. (a) velocity (VCL) at 6 s post-activation across age in years (black: Lake 223, n = 30; grey: Lake 224, n = 30), and (b) decline in sperm swimming velocity with time post-activation in young (pink: 4–10 years, n = 30) and old (blue: 18–37 years, n = 30) fish. Points represent average of technical replicates for each fish, lines represent average among individuals from the same lake and age category. Error bars/bands represent α 1 SE.

### BIOCHEMICIAL PROXY

Relative telomere length in red blood cells was similar in both lakes (Lake 224 – Lake 223: 28 +/− 2308, *t*_56_ = 0.012, *P* = 0.99), and in young and old individuals (old – young: 251+/− 2308, *t*_56.0_ = 0.11, *P* = 0.914; Figure 3c). Relative telomere length in sperm cells was significantly higher in Lake 224 (Lake 224 – Lake 223: 8406 +/− 1673, *t*_57_ = 5.03, *P* <0.0001); however, there was no difference between young and old trout (old – young: −1196 +/− 1673, *t*_57_ = −0.72, *P* = 0.48; Figure 3d).

## 5. DISCUSSSION

As a species, lake trout have evolved under indeterminate growth, and all individuals have this genetic potential. However, realized growth in lake trout varies depending on diet availability, with fish achieving enormous sizes in some lakes, but are stunted in others. We exploited this scenario to make age comparisons among male trout that were not confounded by diet differences. Adult trout in our study lakes have functionally determinate growth due to a simplified foodweb. Despite this, and as predicted by both DSH and DFT, they show little to no senescence in many traits measured here, and while we conclude that overall senescence is negligible in these populations, we argue that there may be negative (reverse) senescence in other populations that are not growth constrained.

Although we have no molecular data to underpin endorsement of DFT, the DSH is clearly supported in our lake trout model. That low adult mortality [relative to juveniles, 6] should be associated with few negative effects of ageing [9, 11] in indeterminate growers [20] paints an incomplete picture of how selection across generations interacts with life history tradeoffs in individuals. Life history theory predicts that consistently low adult mortality *across generations* leads to low reproductive effort in a given year as a means of bet hedging reproductive success across many episodes/years [73]. Under the DSH, through phenotypically plastic allocation of resources, this low reproductive effort would result in high somatic maintenance and thus low senescence *within a generation* (individual). Connecting these concepts for the case of lake trout suggests that due to (1) the lack of adult predation in the growing and spawning environments they evolved under, adult mortality is consistently very low across generations (it is the lowest of any salmonid), (2) resulting in low reproductive effort in a given year (it is the lowest of any salmonid), and (3) through plasticity within-individuals, high somatic maintenance results in no, or limited, senescence, enabling full potential of long life (it is the highest of any salmonid) to hedge reproductive success with environmental stochasticity.

Observed variation in senescent patterns among species [5, 18, 19] suggests contrasting selection pressures as an ultimate cause. Indeterminate growth is predicted [20] to increase selection against senescence when adult individuals experience reduced mortality and increased fertility with age (increasing size). Testing this prediction in wild populations has been challenging due to often confounding variables. Lake trout from our particular populations enable unique opportunities to control such problems, including diet. However, a stable diet likely results in old fish having inferior performance than their inherent potential. Negative senescence is predicted when size at maturity is much smaller than maximum size [20]. In lake trout populations where adults can switch to larger or more energy dense prey as they grow, old fish achieve much larger sizes than young adults [52]. Very large adults would have high sperm quantity (predicted to win under sperm competition = fertility), and high sperm quality (predicted to win under sperm competition = fertility) if under suitable diet (and females would have much more egg production). Such data would not only support negligible senescence, but also show the aptitude for negative (reverse) senescence in this species.

Our study suggests lake trout have at most negligible senescence, with the potential to exhibit negative (reverse) senescence in populations where adults can attain maximum sizes that are much larger than those at maturity, due to prey availability. These data provide support of evolutionary theories of ageing, from rarely studied long-lived indeterminate growing animals in the wild. Our data are unique in that they coalesce information on 1) actuarial senescence using mortality rates from mark-recapture, with 2) measures of phenotypic performance including reproductive senescence. Furthermore, blood and sperm cell telomere lengths did not decline with age, 3) indicating that, at least under the conditions of this study, telomere maintenance through adulthood may in part underpin the lack of apparent senescence. Our age comparisons combine longitudinal (same individuals across decades) with cross-sectional data (difference aged individuals at the same time), which is an infrequent approach. These age assessments are strengthened by the unique characteristics of the study populations that control for confounding variables that are profuse in most natural situations.

If our conclusions are accurate, one can make predictions that should be supported by other data. For instance, (1) old and young males should show equal paternity if tested under sperm competition in the lab, (2) pedigrees of wild populations should show equal average contributions of individual old and young males as fathers in a given year, and (3) laboratory studies should indicate no increase in abnormalities in offspring development from older vs young fathers. (4) If DFT is involved, relative adult to juvenile proxies for cellular hyperfunction, should be correlated with senescence, within and among genera of salmonids in relation to the degree of iteroparity and semelparity, with lake trout at one extreme. (5) Given the unusual insensitivity of relative telomere length to aging in this species, further laboratory and field studies are needed to test if levels of other common molecular markers of senescence in this, and other long-lived ectotherms, may also fail to recapitulate the patterns expected from studies on endotherms or more typical laboratory model organisms. We encourage such studies to be undertaken where possible, along with comparisons across lake trout populations that vary in adult mortality and growth potential.

## ACKNOWLEDGEMENTS

We thank staff of the IISD-ELA for collecting historical information from Lakes 223 and 224, especially K. Mills, S. Chalanchuk and D. Allan. 2017 samples were collected with the assistance of L. Hrenchuk, C. Rogers, L. Hayhurst, A. Milling, C.V. Veen, and M. Fahmy. M. Fahmy also supported the microscope analyses. D. McLennan and K. Jeffries are thanked for assistance with developing and validating the telomere assay.

## Supplementary Methods I – Longitudinal data

### Actuarial senescence

Using all individual adult males with known ages (Lake 223 = 385, Lake 224 = 422), we treated age as a continuous variable and estimated the probability of recapture and the probability of survival (1 - mortality) of each individual adult male in each year (sampled from posterior distributions). As extrinsic adult mortality is low, and young/old adults experience the same conditions, any changes in mortality with age are assumed to be attributed to intrinsic processes.

In order to test for an increase in adult mortality with age, we fitted a Cormack Jolly Seber model (Lebreton *et al.*, 1992) with separate survival (1 – mortality) and capture probabilities as linear regression functions on the logistic scale. We fitted a first order autocorrelation structure for annual random effects on survival and recapture probability, as we expect that factors affecting these parameters (especially survival) are likely to be similar from year to year. The likelihood structure of the basic process model is thus defined as

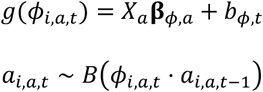

where *a_i,a,t_* is a latent Bernouilli variable indicating whether individual *i* of age *a* is alive at time *i*. An indiviudal will survive from interval *t* – 1 to interval *t* with probability *Φ_i,t_*, only if it was alive at time *t* – 1, i.e., only if *a*_*i,a,t*–1_ = 1. *Φ_i,a,t_* is modelled as a logistic regression with an effect of continuous age on survival contained in β_Φ,a_, and random effects of year, *b_Φ,t_*. The temporal correlation structure of these year effects after the first year is defined by

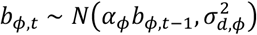

where *α_Φ_* is the first-order autoregressive parameter describing the dependence of survival in one episode on the previous episode and 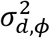 is the variance of disturbances of the autoregressive process. The stationary variance of such a process is 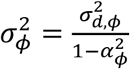, and this variance defines the distribution from which the random effect in the first year is drawn. The observation model takes the form

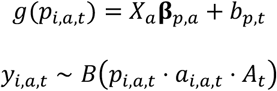

where *y_i,a,t_* is the observation (1 = captured, 0 = not captured) of individual *i* of age *a* at time *t*. Any individual not alive (*a_i,a,t_ =* 0) cannot be observed, and those that are alive may be observed with probability *P_i,a,t_*, provided that sampling was conducted in year *t* (*A_t_* = 1 if sampling was conducted; *A_t_* = 0 otherwise). As in the process (survival) part of the model, the probability of capture of live individuals is modelled as a logistic regression, with separate intercepts for each age, and an autoregressive structure for annual variation. The structure for annual variation is directly analogous to that described above for the survival part of the model.

The model was sampled by Gibbs sampling using jags (Plummer, 2010) in R version 3.6.2. We used diffuse normal priors on all fixed effects and autoregression parameters, and diffuse gamma priors on the precision (inverse of the variance) of the disturbances of the survival and capture parts of the model.

#### Additional parameters

In addition, we tested models that included combinations of a quadratic term for the rate of mortality change and a parameter that varied the minimum mortality rate. However, these models did not converge well and were not numerically stable. This suggests that a quadratic term did not fit our data, and therefore, we did not use these parameters in the final model described above. However, we have left these as options within our scripts for others to see how they were included, alongside the scripts for plots we used to check for convergence.

#### Body condition

Length-based body condition was estimated as a percentage of standard weight (1993). Fish that were recaptured at least 6 times during their adult life were used to determine if condition declined with age, and were analyzed with a mixed effects modelling framework using the *lme4* package (Bates *et al*., 2014) in R. Condition was evaluated as a function of fish age (fixed effect), and repeated measures on the same individuals (modelled as a random slope), and the year sampled (random intercept). Significance of fixed effects was assessed using the Satterthwaite approximation for degrees of freedom with the *lmerTest* package (Kuznetsova *et al*., 2017). Random effects were retained if found to be significant in log-likelihood ratio tests using the *anova()* function in R. Assumptions of normality and homogeneity of variance were verified using model residuals. Only fish from Lake 224 were used in this analysis as exclusion of data prior to 1990 (Mills *et al*., 2000) limited sample sizes in Lake 223.

## Supplemental Methods II – fish used in 2017

**Table S1:**
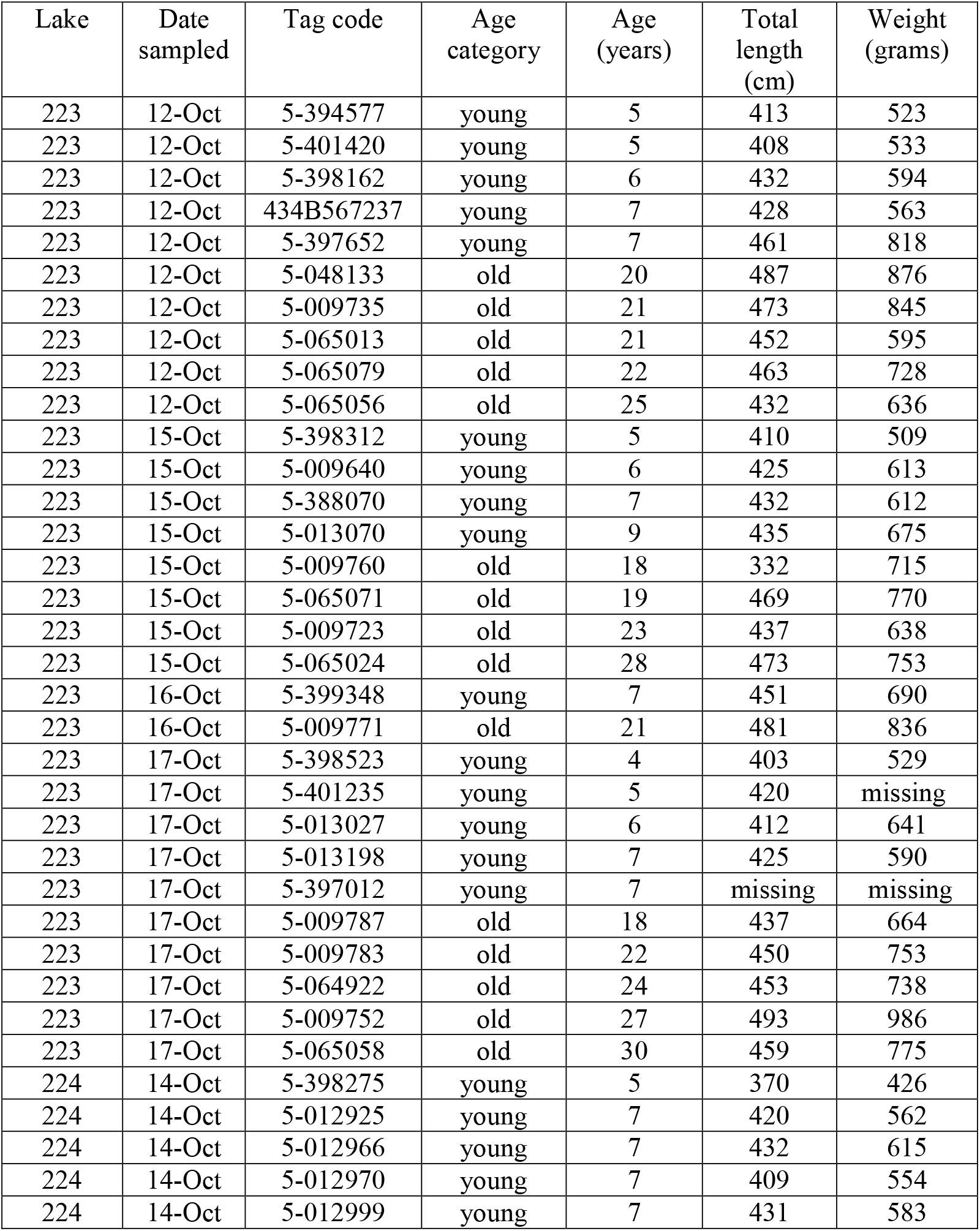
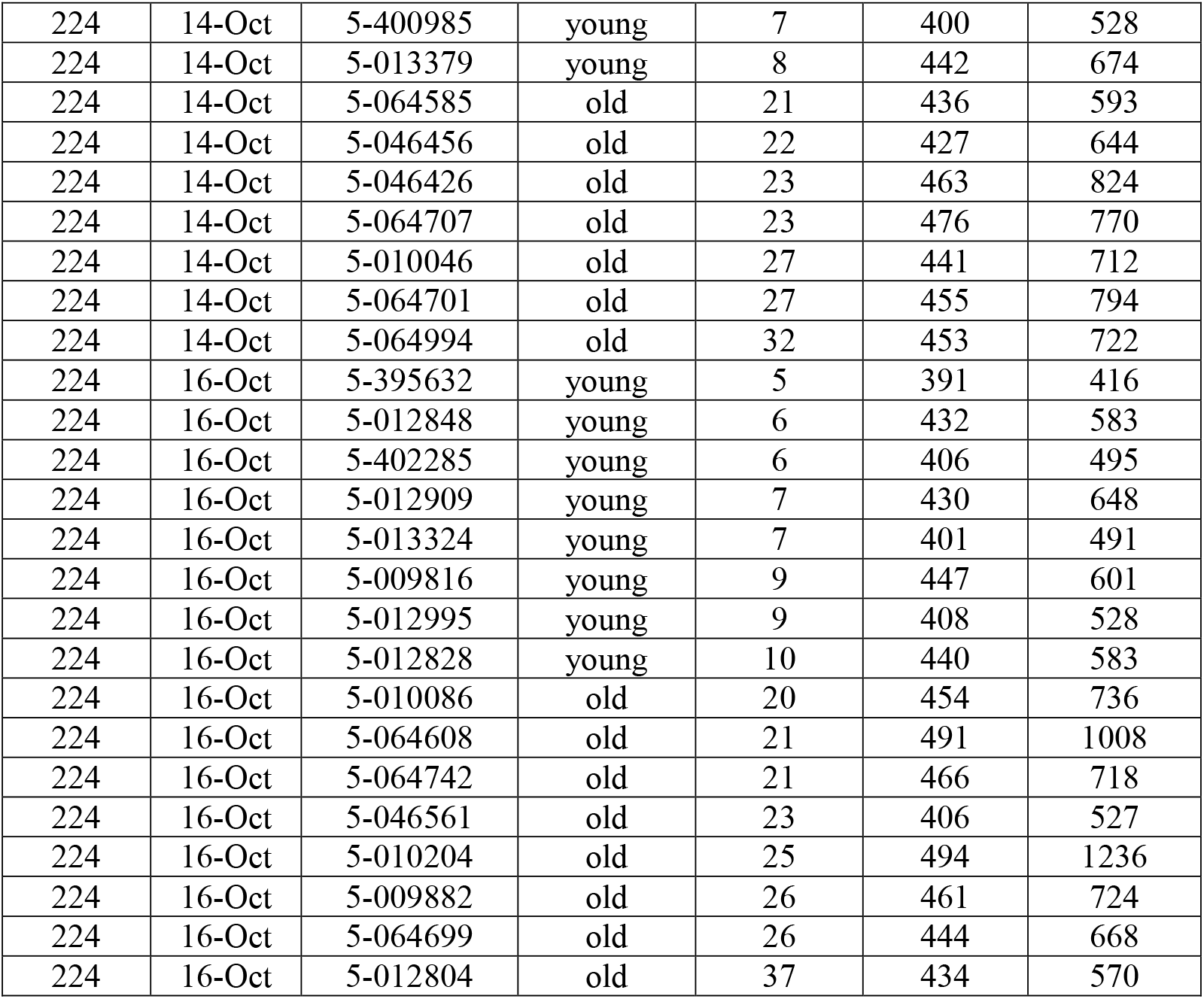
Adult male lake trout sampled from spawning grounds in October 2017. Ages were known in years, and sampled as categories (young = 4–10, old = 18–37). Some data were missing on some fish.

| Lake | Date sampled | Tag code | Age category | Age (years) | Total length (cm) | Weight (grams) |
| --- | --- | --- | --- | --- | --- | --- |
| 223 | 12-Oct | 5-394577 | young | 5 | 413 | 523 |
| 223 | 12-Oct | 5-401420 | young | 5 | 408 | 533 |
| 223 | 12-Oct | 5-398162 | young | 6 | 432 | 594 |
| 223 | 12-Oct | 434B567237 | young | 7 | 428 | 563 |
| 223 | 12-Oct | 5-397652 | young | 7 | 461 | 818 |
| 223 | 12-Oct | 5-048133 | old | 20 | 487 | 876 |
| 223 | 12-Oct | 5-009735 | old | 21 | 473 | 845 |
| 223 | 12-Oct | 5-065013 | old | 21 | 452 | 595 |
| 223 | 12-Oct | 5-065079 | old | 22 | 463 | 728 |
| 223 | 12-Oct | 5-065056 | old | 25 | 432 | 636 |
| 223 | 15-Oct | 5-398312 | young | 5 | 410 | 509 |
| 223 | 15-Oct | 5-009640 | young | 6 | 425 | 613 |
| 223 | 15-Oct | 5-388070 | young | 7 | 432 | 612 |
| 223 | 15-Oct | 5-013070 | young | 9 | 435 | 675 |
| 223 | 15-Oct | 5-009760 | old | 18 | 332 | 715 |
| 223 | 15-Oct | 5-065071 | old | 19 | 469 | 770 |
| 223 | 15-Oct | 5-009723 | old | 23 | 437 | 638 |
| 223 | 15-Oct | 5-065024 | old | 28 | 473 | 753 |
| 223 | 16-Oct | 5-399348 | young | 7 | 451 | 690 |
| 223 | 16-Oct | 5-009771 | old | 21 | 481 | 836 |
| 223 | 17-Oct | 5-398523 | young | 4 | 403 | 529 |
| 223 | 17-Oct | 5-401235 | young | 5 | 420 | missing |
| 223 | 17-Oct | 5-013027 | young | 6 | 412 | 641 |
| 223 | 17-Oct | 5-013198 | young | 7 | 425 | 590 |
| 223 | 17-Oct | 5-397012 | young | 7 | missing | missing |
| 223 | 17-Oct | 5-009787 | old | 18 | 437 | 664 |
| 223 | 17-Oct | 5-009783 | old | 22 | 450 | 753 |
| 223 | 17-Oct | 5-064922 | old | 24 | 453 | 738 |
| 223 | 17-Oct | 5-009752 | old | 27 | 493 | 986 |
| 223 | 17-Oct | 5-065058 | old | 30 | 459 | 775 |
| 224 | 14-Oct | 5-398275 | young | 5 | 370 | 426 |
| 224 | 14-Oct | 5-012925 | young | 7 | 420 | 562 |
| 224 | 14-Oct | 5-012966 | young | 7 | 432 | 615 |
| 224 | 14-Oct | 5-012970 | young | 7 | 409 | 554 |
| 224 | 14-Oct | 5-012999 | young | 7 | 431 | 583 |
| 224 | 14-Oct | 5-400985 | young | 7 | 400 | 528 |
| 224 | 14-Oct | 5-013379 | young | 8 | 442 | 674 |
| 224 | 14-Oct | 5-064585 | old | 21 | 436 | 593 |
| 224 | 14-Oct | 5-046456 | old | 22 | 427 | 644 |
| 224 | 14-Oct | 5-046426 | old | 23 | 463 | 824 |
| 224 | 14-Oct | 5-064707 | old | 23 | 476 | 770 |
| 224 | 14-Oct | 5-010046 | old | 27 | 441 | 712 |
| 224 | 14-Oct | 5-064701 | old | 27 | 455 | 794 |
| 224 | 14-Oct | 5-064994 | old | 32 | 453 | 722 |
| 224 | 16-Oct | 5-395632 | young | 5 | 391 | 416 |
| 224 | 16-Oct | 5-012848 | young | 6 | 432 | 583 |
| 224 | 16-Oct | 5-402285 | young | 6 | 406 | 495 |
| 224 | 16-Oct | 5-012909 | young | 7 | 430 | 648 |
| 224 | 16-Oct | 5-013324 | young | 7 | 401 | 491 |
| 224 | 16-Oct | 5-009816 | young | 9 | 447 | 601 |
| 224 | 16-Oct | 5-012995 | young | 9 | 408 | 528 |
| 224 | 16-Oct | 5-012828 | young | 10 | 440 | 583 |
| 224 | 16-Oct | 5-010086 | old | 20 | 454 | 736 |
| 224 | 16-Oct | 5-064608 | old | 21 | 491 | 1008 |
| 224 | 16-Oct | 5-064742 | old | 21 | 466 | 718 |
| 224 | 16-Oct | 5-046561 | old | 23 | 406 | 527 |
| 224 | 16-Oct | 5-010204 | old | 25 | 494 | 1236 |
| 224 | 16-Oct | 5-009882 | old | 26 | 461 | 724 |
| 224 | 16-Oct | 5-064699 | old | 26 | 444 | 668 |
| 224 | 16-Oct | 5-012804 | old | 37 | 434 | 570 |

## Supplemental Methods III – sperm methods

New sperm activation medium was made each day and contained 79.9% lake water from Lake 239 (site of field station), 20% ovarian fluid, and 0.1% bovine serum albumin, which reduces the likely of sperm sticking to the glass slides (e.g., *Beirão et al*., 2014; Beirão *et al*., 2015). Activation medium and a semen aliquot from each male were kept at 5°C in a temperature-controlled aluminum block next to the microscope. Semen was kept on ice until transfer to the block, and was assessed within 8 hours of stripping. 0.1 μl of semen from a given male was pipetted into the opening of a 2 chamber Cytonix Microtool slide, that was prechilled to 8°C (~ temperature of spawning) using a customized Physitemp TS-4 stage cooling system. This was followed quickly by 3.95 μl of activation media, which mixed with sperm as it filled the slide chamber. We were able to consistently adjust slide position and fine focus within 6 s of sperm/media mixing. Sperm swimming performance was captured at 100 frames per second using a Prosilica GE680 monochrome camera mounted to a Leica DM IL LED inverted microscope with a 20x phase-contrast lens. The entire procedure was repeated four times for each semen sample as a means of technical replication. Videos of swimming sperm were analyzed in 0.5s increments using the Computer Assisted Sperm Analysis (CASA) plugin for ImageJ (Wilson-Leedy and Ingermann, 2007), modified by Purchase and Earle (2012). We used sperm curvilinear swimming velocity (VCL; μm/s) as a metric of sperm quality, as it has been repeatedly shown to be correlated to paternity under sperm competition (e.g., Gage *et al*., 2004; Evans *et al*., 2013; Alonzo *et al*., 2016).

## Supplemental Methods IV – relative telomere length assay

Relative telomere length has been shown to decline with age in several fishes (Rollings *et al*., 2014; Carneiro *et al*., 2016; Hatakeyama *et al*., 2016), including a wild salmonid (McLennan *et al*., 2017), and another long-lived species (Simide *et al*., 2016), although ectotherms do not always show declining telomere length with age (Olsson *et al*., 2018). We measured relative telomere length from DNA recovered from red blood cells and sperm pellets using a qPCR-based approach that produces a telomere repeat (T) to single gene (S) copy number ratio (T/S).

Genomic DNA (gDNA) extractions were performed with 10 μl of RBCs or sperm pellets using a DNeasy Blood and Tissue kit (Qiagen) according to the manufacturer’s directions. The recovered DNA was quantified using a Qubit DNA HS assay kit and Qubit 2.0 Flurometer (Thermo Fisher Scientific) and subsequently diluted to 10 ng/μl. The qPCR-based approach developed by Cawthon (Cawthon, 2002), which produces a telomere repeat (T) to single gene (S) copy number ratio (T/S) for each DNA sample, was used to quantify relative telomere length. Telomere repeats were amplified with the universal primer pair Tel1b and Tel2b from Epel et al. (Epel *et al*., 2004). *Ox* and *FSH* were both used as single copy genes to be able to verify consistency of T/S ratios depending on which single copy gene was targeted (see Supplemental Table I for primer sequences). Primers were designed in Geneious 9.1.8 (Biomatters Ltd.) from publicly available mRNA sequences (Genbank accession numbers HQ656804.1 and HM057170.1 for *Ox* and *FSH*, respectively). qPCRs were performed on separate 384-well plates for each primer pair using the QuantStudio 5 Real-Time PCR System (Thermo Fisher Scientific). Reactions were prepared in triplicate for each sample with 2x PowerUp SYBR Green Master Mix (Thermo Fisher Scientific), 10 ng DNA per reaction, and final concentrations of 800 nM for each primer. Thermocycling conditions for the telomere qPCR were 95°C for 2 min, and 27 cycles of 95°C for 15 s, 56°C for 15 s and 72°C for 60 s. Single copy gene thermocycling conditions were 95°C for 2 min, and 40 cycles of 95°C for 20 s, and 6O°C for 20 s. Both of these qPCR thermocycling programs were followed by default melt curve conditions of 95°C with a ramp rate of 1.6°C/s for 15 s, 60°C with a ramp rate of 1.6/s for 1 min, 95°C with a ramp rate of 0.15°C/s for 15 s. Primer efficiency tests were performed on pooled RBC gDNA from all individual subsamples, which was diluted in a 4-fold serial dilution producing five concentrations ranging 40–0.157 ng/μl. Primer efficiency values ranged from 94-107%. Non-target controls were also performed in triplicate on each plate producing Ct values that matched the background fluorescence values. Relative telomere lengths were calculated as (2^Ct(telomeres)^/2Ct^(single copy gene)^)^−1^ as described by Cawthon (2002).

**Supplemental Table I.**
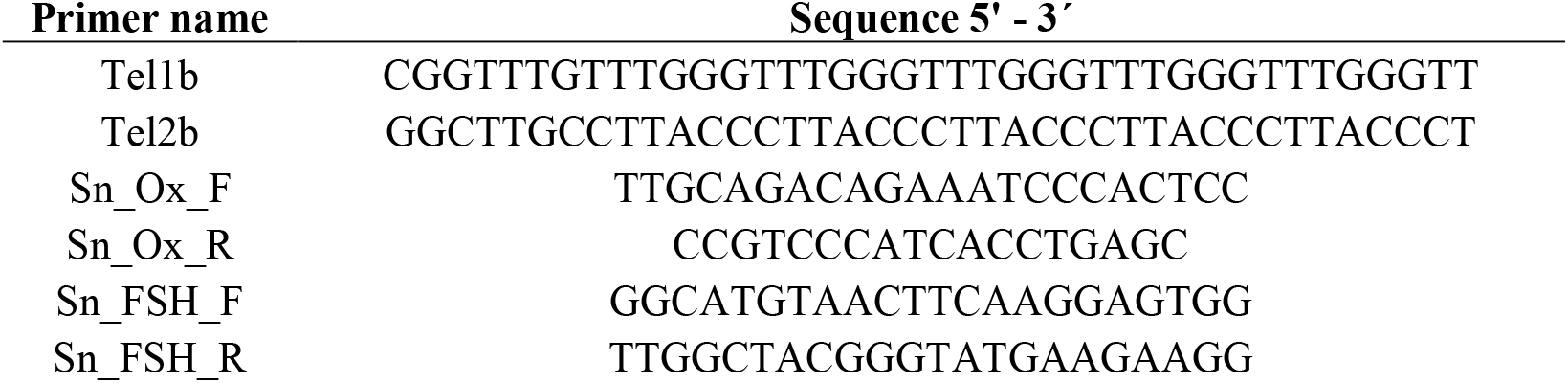
Primers used for relative telomere length assay.

## REFERENCES

1. Nussey D.H., Froy H., Lemaitre J.F., Gaillard J.M., Austad S.N. 2013 Senescence in natural populations of animals: widespread evidence and its implications for bio-gerontology. Ageing Res Rev 12(1), 214–225. (doi:10.1016/j.arr.2012.07.004).

2. Bernard C., Compagnoni A., Salguero-Gómez R. 2020 Testing Finch’s hypothesis: the role of organismal modularity on the escape from actuarial senescence. Functional Ecology 34(1), 88–106. (doi:10.1111/1365-2435.13486).

3. Jones O.R., Scheuerlein A., Salguero-Gomez R., Camarda C.G., Schaible R., Casper B.B., Dahlgren J.P., Ehrlen J., Garcia M.B., Menges E.S., et al. 2014 Diversity of ageing across the tree of life. Nature 505(7482), 169–173. (doi:10.1038/nature12789).

4. Baudisch A., Stott I. 2019 A pace and shape perspective on fertility. Methods in Ecology and Evolution 10(11), 1941–1951. (doi:10.1111/2041-210x.13289).

5. Baudisch A., Vaupel J.W. 2012 Getting to the root of aging. Science 338(6107), 618–619. (doi:10.1126/science.1226467).

6. Moorad J., Promislow D., Silvertown J. 2019 Evolutionary ecology of senescence and a reassessment of Williams’‘extrinsic mortality’ hypothesis. Trends Ecol Evol 34(6), 519–530. (doi:10.1016/j.tree.2019.02.006).

7. Roper M., Capdevila P., Salguero-Gómez R. 2021 Senescence: why and where selection gradients might not decline with age. Proceedings of the Royal Society B: Biological Sciences 288(1955), 20210851. (doi:doi:10.1098/rspb.2021.0851).

8. Maklakov A.A., Chapman T. 2019 Evolution of ageing as a tangle of trade-offs: energy versus function. Proc Biol Sci 286(1911), 20191604. (doi:10.1098/rspb.2019.1604).

9. Williams G.C. 1957 Pleiotropy, natural selection, and the evolution of senescence. Evolution 11(4), 398–411. (doi:10.1111/j.1558-5646.1957.tb02911.x).

10. da Silva J. 2018 Reports of the death of extrinsic mortality moulding senescence have been greatly exaggerated. Evol Biol 45(2), 140–143. (doi:10.1007/s11692-018-9446-y).

11. Danko M.J., Burger O., Argasinski K., Kozlowski J. 2018 Extrinsic mortality can shape life-history traits, including senescence. Evol Biol 45(4), 395–404. (doi:10.1007/s11692-018-9458-7).

12. Medawar P.B. 1952 An unsolved problem of biology. London, H.K. Lewis.

13. Kirkwood T.B.L. 1977 Evolution of aging. Nature 270(5635), 301–304. (doi:10.1038/270301a0).

14. Blagosklonny M.V. 2006 Aging and immortality: quasi-programmed senescence and its pharmacologic inhibition. Cell Cycle 5(18), 2087–2102. (doi:10.4161/cc.5.18.3288).

15. Cohen A.A., Coste C.F.D., Li X.-Y., Bourg S., Pavard S., Gaillard J.-M. 2020 Are trade-offs really the key drivers of ageing and life span? Functional Ecology 34(1), 153–166. (doi:10.1111/1365-2435.13444).

16. Gaillard J.M., Lemaître J.F., Fox C. 2020 An integrative view of senescence in nature. Functional Ecology 34(1), 4–16. (doi:10.1111/1365-2435.13506).

17. Lind M.I., Ravindran S., Sekajova Z., Carlsson H., Hinas A., Maklakov A.A. 2019 Experimentally reduced insulin/IGF-1 signaling in adulthood extends lifespan of parents and improves Darwinian fitness of their offspring. Evol Lett 3(2), 207–216. (doi:10.1002/evl3.108).

18. Finch C.E. 1998 Variations in senescence and longevity include the possibility of negligible senescence. Journals of Gerontology Series a-Biological Sciences and Medical Sciences 53(4), B235–B239. (doi:10.1093/gerona/53A.4.B235).

19. Finch C.E. 2009 Update on slow aging and negligible senescence--a mini-review. Gerontology 55(3), 307–313. (doi:10.1159/000215589).

20. Vaupel J.W., Baudisch A., Dolling M., Roach D.A., Gampe J. 2004 The case for negative senescence. Theor Popul Biol 65(4), 339–351. (doi:10.1016/j.tpb.2003.12.003).

21. Fletcher Q.E., Selman C. 2015 Aging in the wild: insights from free-living and non-model organisms. Exp Gerontol 71, 1–3. (doi:10.1016/j.exger.2015.09.015).

22. Olsson M., Shine R. 1996 Does reproductive success increase with age or with size in species with indeterminate growth? A case study using sand lizards *(Lacerta agilis)*. Oecologia 105(2), 175–178. (doi:10.1007/bf00328543).

23. Roach D.A., Smith E.F., Gaillard J.-M. 2020 Life-history trade-offs and senescence in plants. Functional Ecology 34(1), 17–25. (doi:10.1111/1365-2435.13461).

24. Hoekstra L.A., Schwartz T.S., Sparkman A.M., Miller D.A.W., Bronikowski A.M., Lemaître J.F. 2020 The untapped potential of reptile biodiversity for understanding how and why animals age. Functional Ecology 34(1), 38–54. (doi:10.1111/1365-2435.13450).

25. Bonduriansky R., Brassil C.E. 2002 Rapid and costly ageing in wild male flies. Nature 420(6914), 377–377. (doi:10.1038/420377a).

26. Monaghan P., Charmantier A., Nussey D.H., Ricklefs R.E. 2008 The evolutionary ecology of senescence. Functional Ecology 22(3), 371–378. (doi:10.1111/j.1365-2435.2008.01418.x).

27. Johnson S.L., Zellhuber-McMillan S., Gillum J., Dunleavy J., Evans J.P., Nakagawa S., Gemmell N.J. 2018 Evidence that fertility trades off with early offspring fitness as males age. Proceedings of the Royal Society B-Biological Sciences 285(1871). (doi:20172174 10.1098/rspb.2017.2174).

28. Roach D.A., Carey J.R. 2014 Population biology of aging in the wild. Annual Review of Ecology, Evolution, and Systematics 45(1), 421–443. (doi:10.1146/annurev-ecolsys-120213-091730).

29. Johnson S.L., Gemmell N.J. 2012 Are old males still good males and can females tell the difference? Do hidden advantages of mating with old males off-set costs related to fertility, or are we missing something else? Bioessays 34(7), 609–619. (doi:10.1002/bies.201100157).

30. Angelier F., Weimerskirch H., Barbraud C., Chastel O., Hopkins W. 2019 Is telomere length a molecular marker of individual quality? Insights from a long-lived bird. Functional Ecology 33(6), 1076–1087. (doi:10.1111/1365-2435.13307).

31. Monaghan P., Metcalfe N.B. 2019 The deteriorating soma and the indispensable germline: gamete senescence and offspring fitness. Proc Biol Sci 286(1917), 20192187. (doi:10.1098/rspb.2019.2187).

32. Lemaitre J.F., Berger V., Bonenfant C., Douhard M., Gamelon M., Plard F., Gaillard J.M. 2015 Early-late life trade-offs and the evolution of ageing in the wild. Proc Biol Sci 282(1806), 20150209. (doi:10.1098/rspb.2015.0209).

33. Lemaitre J.F., Gaillard J.M. 2017 Reproductive senescence: new perspectives in the wild. Biol Rev Camb Philos Soc 92(4), 2182–2199. (doi:10.1111/brv.12328).

34. Graves B.M. 2007 Sexual selection effects on the evolution of senescence. Evolutionary Ecology 21(5), 663–668. (doi:10.1007/s10682-006-9144-6).

35. Bartosch-Harlid A., Berlin S., Smith N.G.C., Moller A.P., Ellegren H. 2003 Life history and the male mutation bias. Evolution 57(10), 2398–2406. (doi:10.1554/03-036).

36. Beirne C., Delahay R., Young A. 2015 Sex differences in senescence: the role of intra-sexual competition in early adulthood. Proc Biol Sci 282(1811). (doi:10.1098/rspb.2015.1086).

37. Bonduriansky R., Maklakov A., Zajitschek F., Brooks R. 2008 Sexual selection, sexual conflict and the evolution of ageing and life span. Functional Ecology 22(3), 443–453. (doi:10.1111/j.1365-2435.2008.01417.x).

38. Lemaitre J.F., Gaillard J.M., Pemberton J.M., Clutton-Brock T.H., Nussey D.H. 2014 Early life expenditure in sexual competition is associated with increased reproductive senescence in male red deer. Proc Biol Sci 281(1792). (doi:10.1098/rspb.2014.0792).

39. Metcalf C.J.E., Roth O., Graham A.L., Lemaître J.-F. 2020 Why leveraging sex differences in immune trade-offs may illuminate the evolution of senescence. Functional Ecology 34(1), 129–140. (doi:10.1111/1365-2435.13458).

40. Grunst M.L., Grunst A.S., Formica V.A., Korody M.L., Betuel A.M., Barcelo-Serra M., Gonser R.A., Tuttle E.M. 2018 Actuarial senescence in a dimorphic bird: different rates of ageing in morphs with discrete reproductive strategies. Proceedings of the Royal Society B-Biological Sciences 285(1892). (doi:20182053 10.1098/rspb.2018.2053).

41. Maklakov A.A., Immler S. 2016 The expensive germline and the evolution of ageing. Current Biology 26(13), R577–R586. (doi:10.1016/j.cub.2016.04.012).

42. Pizzari T., Dean R., Pacey A., Moore H., Bonsall M.B. 2008 The evolutionary ecology of pre- and post-meiotic sperm senescence. Trends Ecol Evol 23(3), 131–140. (doi:10.1016/j.tree.2007.12.003).

43. Reinhardt K., Turnell B. 2019 Sperm ageing: a complex business. Functional Ecology 33(7), 1188–1189. (doi:10.1111/1365-2435.13350).

44. Purchase C.F., Evans J.P., Roncal J. 2021 Intergrating natural and sexual selection across the biphasic life cycle. EcoEvoRxiv. (doi:https://doi.org/10.32942/osf.io/eu3am).

45. Vega-Trejo R., Fox R.J., Iglesias-Carrasco M., Head M.L., Jennions M.D., Priest N. 2019 The effects of male age, sperm age and mating history on ejaculate senescence. Functional Ecology 33(7), 1267–1279. (doi:10.1111/1365-2435.13305).

46. Crow J.F. 2000 The origins patterns and implications of human spontaneous mutation. Nature Reviews Genetics 1(1), 40–47. (doi:10.1038/35049558).

47. Gasparini C., Devigili A., Pilastro A. 2019 Sexual selection and ageing: interplay between pre- and post-copulatory traits senescence in the guppy. Proceedings of the Royal Society B-Biological Sciences 286(1897). (doi:20182873 10.1098/rspb.2018.2873).

48. Aich U., Head M.L., Fox R.J., Jennions M.D. 2021 Male age alone predicts paternity success under sperm competition when effects of age and past mating effort are experimentally separated. Proceedings of the Royal Society B: Biological Sciences 288(1955), 20210979. (doi:doi:10.1098/rspb.2021.0979).

49. Behnke R.J. 2002 Trout and salmon of North America. New York, The Free Press; 359 p.

50. Esteve M., McLennan D.A., Gunn J.M. 2007 Lake trout *(Salvelinus namaycush)* spawning behaviour: the evolution of a new female strategy. Environmental Biology of Fishes 83(1), 69–76. (doi:10.1007/s10641-007-9272-z).

51. Parker G.A. 1990 Sperm competition games - raffles and roles. Proceedings of the Royal Society B-Biological Sciences 242(1304), 120–126. (doi:10.1098/rspb.1990.0114).

52. Cruz-Font L., Shuter B.J., Blanchfield P.J., Minns C.K., Rennie M.D. 2019 Life at the top: lake ecotype influences the foraging pattern, metabolic costs and life history of an apex fish predator. Journal of Animal Ecology 88(5), 702–716. (doi:10.1111/1365-2656.12956).

53. Kennedy P.J., Bartley T.J., Gillis D.M., McCann K.S., Rennie M.D. 2018 Offshore prey densities facilitate similar life history and behavioral patterns in two distinct aquatic apex predators, northern pike and lake trout. Transactions of the American Fisheries Society 147(5), 972–995. (doi:10.1002/tafs.10090).

54. Purchase C.F., Collins N.C., Shuter B.J. 2005 Sensitivity of maximum sustainable harvest rates to intra-specific life history variability of lake trout *(Salvelinus namaycush)* and walleye *(Sander vitreus)*. Fisheries Research 72(2-3), 141–148. (doi:10.1016/j.fishres.2004.11.006).

55. Kirkwood T.B., Melov S. 2011 On the programmed/non-programmed nature of ageing within the life history. Curr Biol 21(18), R701–707. (doi:10.1016/j.cub.2011.07.020).

56. Campana S.E., Casselman J.M., Jones C.M. 2008 Bomb radiocarbon chronologies in the Arctic, with implications for the age validation of lake trout *(Salvelinus namaycush)* and other Arctic species. Canadian Journal of Fisheries and Aquatic Sciences 65(4), 733–743. (doi:10.1139/f08-012).

57. Mills K.H., Chalanchuk S.M., Allan D.J. 2000 Recovery of fish populations in Lake 223 from experimental acidification. Canadian Journal of Fisheries and Aquatic Sciences 57(1), 192–204. (doi:10.1139/cjfas-57-1-192).

58. Butts I.A.E., Hilmarsdóttir G.S., Zadmajid V., Gallego V., Støttrup J.G., Jacobsen C., Krüger-Johnsen M., Politis S.N., Asturiano J.F., Holst L.K., et al. 2020 Dietary amino acids impact sperm performance traits for a catadromous fish, *Anguilla anguilla* reared in captivity. Aquaculture 518, 734602. (doi:10.1016/j.aquaculture.2019.734602).

59. Blackwell B.G., Brown M.L., Willis D.W. 2000 Relative weight (Wr) status and current use in fisheries assessment and management. Reviews in Fisheries Science 8(1), 1–44. (doi:10.1080/10641260091129161).

60. Hartman K.J., Margraf F.J. 2006 Relationships among condition indicies, feeding and growth of wallye in Lake Erie. Fisheries Management and Ecology 13, 121–130.

61. Carneiro M.C., de Castro I.P., Ferreira M.G. 2016 Telomeres in aging and disease: lessons from zebrafish. Dis Model Mech 9(7), 737–748. (doi:10.1242/dmm.025130).

62. Hatakeyama H., Yamazaki H., Nakamura K.-I., Izumiyama-Shimomura N., Aida J., Suzuki H., Tsuchida S., Matsuura M., Takubo K., Ishikawa N. 2016 Telomere attrition and restoration in the normal teleost *Oryzias latipes* are linked to growth rate and telomerase activitiy at each life stage. Aging 8, 62–75.

63. Rollings N., Miller E., Olsson M. 2014 Telomeric attrition with age and temperature in Eastern mosquitofish *(Gambusia holbrooki)*. Naturwissenschaften 101(3), 241–244. (doi:10.1007/s00114-014-1142-x).

64. Mills K.H., Beamish R.J. 1980 Comparisons of fin-ray and scale age-determinations for lake whitefish *(Coregonus clupeaformis)* and their implications for estimates of growth and annual survival. Canadian Journal of Fisheries and Aquatic Sciences 37(3), 534–544. (doi:10.1139/f80-068).

65. Piccolo J.J., Hubert W.A., Whaley R.A. 1993 Standard weight equation for lake trout. North American Journal of Fisheries Management 13(2), 401–404. (doi:10.1577/1548-8675(1993)013<0401:sweflt>2.3.co;2).

66. Rennie M.D., Kennedy P.J., Mills K.H., Rodgers C.M.C., Charles C., Hrenchuk L.E., Chalanchuk S., Blanchfield P.J., Paterson M.J., Podemski C.L. 2019 Impacts of freshwater aquaculture on fish communities: a whole-ecosystem experimental approach. Freshwater Biology 64(5), 870–885. (doi:10.1111/fwb.13269).

67. Purchase C.F., Rooke A.C. 2020 Freezing ovarian fluid does not alter how it affects fish sperm swimming performance: creating a cryptic female choice ‘spice rack’ for use in split-ejaculate experimentation. Journal of Fish Biology 96(3), 693–699. (doi:10.1111/jfb.14263).

68. Purchase C.F., Moreau D.T. 2012 Stressful environments induce novel phenotypic variation: hierarchical reaction norms for sperm performance of a pervasive invader. Ecol Evol 2(10), 2567–2576. (doi:10.1002/ece3.364).

69. Purchase C.F., Butts I.A.E., Alonso-Fernández A., Trippel E.A. 2010 Thermal reaction norms in sperm performance of Atlantic cod *(Gadus morhua)*. Canadian Journal of Fisheries and Aquatic Sciences 67(3), 498–510. (doi:10.1139/f10-001).

70. Johnson K., Butts I.A.E., Wilson C.C., Pitcher T.E. 2013 Sperm quality of hatchery-reared lake trout throughout the spawning season. North American Journal of Aquaculture 75(1), 102–108. (doi:10.1080/15222055.2012.711277).

71. Purchase C.F., Earle P.T. 2012 Modifications to the ImageJ computer assisted sperm analysis plugin greatly improve efficiency and fundamentally alter the scope of attainable data. Journal of Applied Ichthyology 28(6), 1013–1016. (doi:10.1111/jai.12070).

72. Gage M.J.G., Macfarlane C.P., Yeates S., Ward R.G., Searle J.B., Parker G.A. 2004 Spermatozoal traits and sperm competition in Atlantic salmon. Current Biology 14(1), 44–47. (doi:10.1016/j.cub.2003.12.028).

73. Roff D.A. 2002 Life history evolution. Sunderland, Sinauer; 256 p.

## References

Bates, D., Maechler, M., Bolker, B. & Walker, S. (2014). Lme4: Linear mixed-effects models using eigen and S4., p. Retrieved from https://cran.rproject.org/web/packages/lme4.

Kuznetsova, A., Brockhoff, P. B. & Christensen, R. H. B. (2017). lmerTest Package: tests in Linear Mixed Effects Models. Journal of Statistical Software 82.

Lebreton, J.-D., Burnham, K. P., Clobert, J. & Anderson, D. R. (1992). Modeling survival and testing biological hypotheses using marked animals: a unified approach with case studies. Ecological Monographs 62, 67–118.

Mills, K. H., Chalanchuk, S. M. & Allan, D. J. (2000). Recovery of fish populations in Lake 223 from experimental acidification. Canadian Journal of Fisheries and Aquatic Sciences 57, 192–204.

Piccolo, J. J., Hubert, W. A. & Whaley, R. A. (1993). Standard weight equation for lake trout. North American Journal of Fisheries Management 13, 401–404.

Plummer, M. (2010). Jags version 2.2.0 user manual.

## References

Alonzo, S. H., Stiver, K. A. & Marsh-Rollo, S. E. (2016). Ovarian fluid allows directional cryptic female choice despite external fertilization. Nature communications 7, 12452.

Beirão, J., Purchase, C. F., Wringe, B. F. & Fleming, I. A. (2014). Sperm plasticity to seawater temperatures in Atlantic cod *Gadus morhua* is affected more by population origin than individual environmental exposure. Marine Ecology Progress Series 495, 263–274.

Beirão, J., Purchase, C. F., Wringe, B. F. & Fleming, I. A. (2015). Inter-population ovarian fluid variation differentially modulates sperm motility in Atlantic cod *Gadus morhua*. Journal of Fish Biology 87, 54–68.

Evans, J. P., Rosengrave, P., Gasparini, C. & Gemmell, N. J. (2013). Delineating the roles of males and females in sperm competition. Proceedings. Biological sciences 280, 20132047.

Gage, M. J. G., Macfarlane, C. P., Yeates, S., Ward, R. G., Searle, J. B. & Parker, G. A. (2004). Spermatozoal traits and sperm competition in Atlantic salmon. Current Biology 14, 44–47.

Purchase, C. F. & Earle, P. T. (2012). Modifications to the ImageJ computer assisted sperm analysis plugin greatly improve efficiency and fundamentally alter the scope of attainable data. Journal of Applied Ichthyology 28, 1013–1016.

Wilson-Leedy, J. G. & Ingermann, R. L. (2007). Development of a novel CASA system based on open source software for characterization of zebrafish sperm motility parameters. Theriogenology 67, 661–672.

## References

Carneiro, M. C., de Castro, I. P. & Ferreira, M. G. (2016). Telomeres in aging and disease: lessons from zebrafish. Disease models & mechanisms 9, 737–748.

Cawthon, R. M. (2002). Telomere measurement by quantitative PCR. Nucleic Acids Research 30, e47.

Epel, E. S., Blackburn, E. H., Lin, J., Dhabhar, F. S., Adler, N. E., Morrow, J. D. & Cawthon, R. M. (2004). Accelerated telomere shortening in response to life stress. Proceedings of the National Academy of Sciences of the United States of America 101, 17312–17315.

Hatakeyama, H., Yamazaki, H., Nakamura, K.-I., Izumiyama-Shimomura, N., Aida, J., Suzuki, H., Tsuchida, S., Matsuura, M., Takubo, K. & Ishikawa, N. (2016). Telomere attrition and restoration in the normal teleost *Oryzias latipes* are linked to growth rate and telomerase activitiy at each life stage. Aging 8, 62–75.

McLennan, D., Armstrong, J. D., Stewart, D. C., McKelvey, S., Boner, W., Monaghan, P., Metcalfe, N. B. & Williams, T. (2017). Shorter juvenile telomere length is associated with higher survival to spawning in migratory Atlantic salmon. Functional Ecology 31, 2070–2079.

Olsson, M., Wapstra, E. & Friesen, C. (2018). Ectothermic telomeres: it’s time they came in from the cold. Philosophical transactions of the Royal Society of London. Series B, Biological sciences 373.

Rollings, N., Miller, E. & Olsson, M. (2014). Telomeric attrition with age and temperature in Eastern mosquitofish *(Gambusia holbrooki)*. Die Naturwissenschaften 101, 241–244.

Simide, R., Angelier, F., Gaillard, S. & Stier, A. (2016). Age and heat stress as determinants of telomere length in a long-lived fish, the Siberian sturgeon. Physiological and biochemical zoology: PBZ 89, 441–447.

